# Specialised root hair cells facilitate rhizobial infection

**DOI:** 10.64898/2026.04.10.717526

**Authors:** Manuel Frank, Huijun Liu, Lavinia Ionna Fechete, Jule Salfeld, Fabian van Beveren, Emma Kleister Sørensen Birkeskov, Henriette Rübsam, Nikolaj Birkebæk Abel, Marcin Nadzieja, Mengyu Lei, Pierre-Marc Delaux, Kasper Røjkjær Andersen, Thomas Ott, Jens Stougaard, Dugald Reid, Stig Uggerhøj Andersen

## Abstract

Legumes establish symbiotic partnerships with soil bacteria that convert atmospheric nitrogen into plant-available forms. Symbiotic bacteria enter through root hairs following recognition by cell surface receptors that help identify compatible symbionts. However, many root hairs express these receptors, and it has long remained unclear why only a small fraction become infected. Here, we use single-cell transcriptomics to show that legumes pre-specify a rare root hair population for infection before bacterial contact. These susceptible root hairs represent less than one percent of the total, express infection-associated genes prior to encountering symbionts and are conserved in distantly related legumes. Their abundance is regulated by the hormone ethylene and correlates with infection capacity. Our findings reveal that root hair cells do not respond uniformly to symbionts but are instead transcriptionally specialised in advance to control infection entry points. This pre-specification provides a mechanism to balance symbiotic benefits against pathogen infection risks and may exemplify a more general strategy used by multicellular hosts to spatially restrict microbial access.

## Main

Many mutualistic interactions rely on hosts accommodating beneficial partners in a spatially confined set of cells rather than throughout their tissues. In intimate symbioses that require intracellular colonization, controlling which host cells can be colonised strongly influences both the efficiency and the ultimate outcome of the association ^1,2^. Yet for most symbioses, it is unknown whether symbiont-hosting cells are selected stochastically from a uniform population or are instead developmentally pre-specified before partner contact occurs.

The symbiosis between legumes and nitrogen-fixing rhizobia presents a striking case of spatial restriction. Root hairs (RHs) in the susceptible zone serve as the entry point for rhizobial infection, and most of these cells express plasma membrane-localised receptors capable of perceiving bacterial lipochitooligosaccharide signals (Nod factors, NFs) ^3–5^. Perception of NFs secreted by compatible rhizobia triggers rapid calcium oscillations, transcriptional reprogramming, and coordinated developmental responses that culminate in nodule primordia formation and bacterial infection ^6,7^. Via these signalling pathways, external microbial cues activate a latent cellular potential, orchestrating infection threads (ITs), cell-to-cell bacterial passage, and the differentiation of bacteroid-containing symbiosomes. From this perspective, host cell symbiotic competence arises as a reaction to symbiont perception and the subsequent molecular dialogue.

However, this canonical view conflicts with a long-standing quantitative discrepancy between how many cells can perceive symbionts and how many are actually infected. Although most RHs in the susceptible zone are theoretically able to perceive rhizobial signals via the NF receptors (NFRs), only between 0.3 and 5 % of these cells are actually colonised ^8^. This disparity between perception capacity and infection frequency has remained unexplained. Several mechanisms shape infection outcomes after rhizobial perception. Nitrogen availability perceived through NLP transcription factors ^9,10^ and autoregulation of nodulation ^11,12^ suppress nodulation systemically. Phytohormones including ethylene, cytokinin, and auxin modulate root susceptibility ^12^ through transcriptional regulators such as NIN, NSP1, and NSP2 ^13–15^. Below infected RHs, cortical cells undergo chromatin remodelling and cytoplasmic reorganisation via calcium signalling, priming them for IT passage ^16,17^. As all of these processes occur after rhizobial perception they do not explain why particular RHs, rather than their neighbours, become infected in the first place. Moreover, the restriction of infection to the susceptible zone near the root tip suggests that competence is developmentally defined ^18^, but whether all RHs within this zone possess equal infection potential, or whether a subpopulation is uniquely competent, has not been determined.

Here, we challenge the prevailing reactive model by identifying a rare, ethylene-regulated RH population that, already prior to bacterial contact, transcriptionally resembles infected cells and is pre-disposed for rhizobium infection.

### Susceptible RHs express infection-related genes prior to rhizobia contact

Although many RHs express *NFR*s and are likely capable of perceiving NFs ^19^, only a small fraction, fewer than five percent, are actually colonised by rhizobia ^8^. We hypothesised that some uninoculated RHs might already resemble infected cells at the transcriptional level, representing a population susceptible to infection (**Figure 1a**).

**Figure 1.**
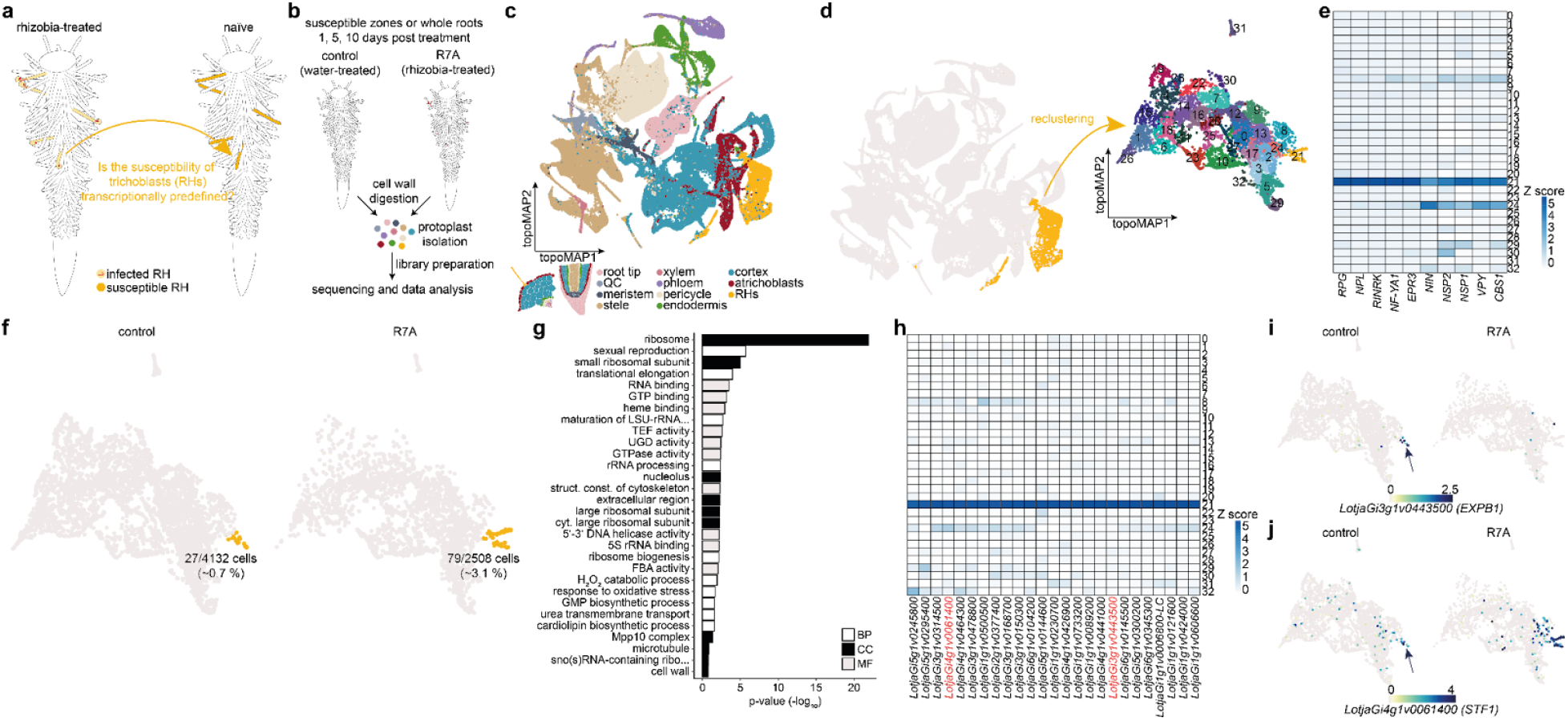
Specialised RHs express rhizobia-induced genes before inoculation. **a**) Central hypothesis: Successfully infected RHs are transcriptionally predefined by susceptibility gene expression prior to rhizobia perception in naive plants. **b**) Summary of experimental procedure used for scRNA-seq data generation. In multiple independent experiments, wild-type plants were treated with water (control) or inoculated with rhizobium *M. loti* R7A (R7A) and cultivated for 1, 5 or 10 days. After incubation, susceptible zones or whole roots were harvested, cell walls digested, and protoplasts isolated. Protoplasts were then used for subsequent library preparation. **c**) topoMAP comprising all wild-type root cells from all experiments. QC: quiescent center. **d**) RHs (gold) were isolated and reclustered resulting in 33 subclusters. **e**) Expression of known infection-related genes in RH subclusters. **f**) RH cells split by treatment. Subcluster 21 containing susceptible and rhizobia-responsive RHs is colored in gold while all other RH cells are grey. Percentage indicates the proportion of subcluster 21 cells compared to all RHs in one treatment. **g**) GO term analysis of control cluster 21 marker genes. BP: biological process, CC: cell compartment, MF: molecular function. GO term analysis lists can be found in Source Data “Supp data S2 - GO term analysis”. **h**) Expression patterns of the top 25 subcluster 21 control marker genes in RH subclusters under control conditions. Gene IDs in red indicate genes depicted in i-j. **i**) Normalised *EXPANSIN B1* (*EXPB1*) expression in RHs. **j**) Normalised *STICKY FINGERS1* (*STF1*) expression in RHs. Arrows indicate susceptible RHs.

Based on single-cell RNA sequencing (scRNA-seq) data, cells can be clustered based on transcriptome similarity. Thus, we reasoned that susceptible RHs could cluster with infected RHs, and that susceptibility marker genes might promote infection. To maximize statistical power for identifying rare cell populations and overcome inherent technical noise in scRNA-seq ^20^, we integrated five independent *Lotus japonicus* scRNA-seq experiments, encompassing wild-type roots harvested at 1, 5, and 10 days post treatment with water (control) or inoculation with *Mesorhizobium loti* R7A ^21,22^ (**Figure 1b, Supplemental Figure S1**). The integrated dataset included 115841 high-quality cells, each with approximately 1,000 detected features (**Figure 1c, Supplemental Figure S1**). In our TopOMetry-clustered ^23^ object, we identified 6640 cells (5.73 % of total cell number) as RH cells based on previously published annotations ^21^ and re-clustered them into 33 subclusters (**Figure 1d, Supplemental Figure S1, Source Data “Supp data S1 - sc RH gene lists”**). Subcluster 21 was identified as infection-related, as the majority of its cells expressed known infection marker genes like *CBS1* ^24^, *NSP2* ^14,25^ and *EXPA1* ^26^ (**Figure 1e, Source Data “Supp data S1 - sc RH gene lists” sheet “wt c21 SRH”**). Compared to these highly specific infection marker genes, *NFR1* and *NFR5* were expressed in about half of all RHs before inoculation and were downregulated after rhizobia perception (**Supplemental Figure S2**). Subcluster 21 contained 106 cells, of which 27 originated from uninoculated control samples, representing 0.7 % of all control RHs (n = 4132 control RH cells; **Figure 1f**). We designated these cells ‘susceptible RHs’ based on their transcriptional resemblance to infected cells and identified 611 significantly enriched susceptibility marker genes of which 163 were present in a minimum of 15 % of susceptible RHs and absent in minimum 85 % of all other RHs (**Source Data “Supp data S1 - sc RH gene lists” sheet “wt c21 SRH”**). Performing gene ontology term enrichment ^27^ for all 611 marker genes revealed an overrepresentation of genes related to the cell wall, extracellular space or H_2_O_2_ catabolism (**Figure 1g, Source Data “Supp data S2 - GO term analysis”**), consistent with cellular functions required for and cellular compartment involved in bacterial accommodation. The top 25 genes showed high specificity for cluster 21 but were also moderately expressed in neighboring clusters 8 and 24 (**Figure 1h**), suggesting a transcriptional gradient of infection competence. Comparing control and inoculated samples, we identified 28 upregulated susceptible RH genes and 10 downregulated susceptible RH genes (**Source Data “Supp data S1 - sc RH gene lists” sheet “wt R7Avscontrol c21”**). This dynamic response demonstrates that susceptible RHs are not merely transcriptionally ‘frozen’ in an infection-like state but modulate expression upon bacterial perception. Upregulated genes included known infection mediators such as *NPL* ^28,29^, *SYMRKL1* ^21^ and *NF-YA1* ^30^, suggesting our approach successfully captured *bona fide* infection determinants. For detailed functional validation, we selected two genes representing distinct regulatory patterns. The first one was the rhizobia-downregulated susceptibility gene *LotjaGi3g1v0443500* encoding an EXPANSIN B1 (EXPB1), a cell wall-loosening protein (**Figure 1i**). The second was a rhizobia-induced susceptibility gene *LotjaGi4g1v0061400* encoding an E6-like protein, which we named STICKY FINGERS1 (STF1, **Figure 1j**). We generated promoter-tYFPnls reporter constructs and studied the expression in transgenic hairy roots. tYFP signal was present in RH nuclei before rhizobia application, and also in RHs of inoculated hairy roots (**Supplemental Figures S3 - S5**).

### *STICKY FINGERS1* is required for rhizobial infection

To investigate the functional importance of a susceptibility marker gene, we examined *STF1* further. *STF1* encodes an E6-like protein, whose orthologues in Arabidopsis, tomato and cotton are involved in fiber elongation, pollen-stigma interactions and fruit ripening ^31–33^. It encodes for 297 amino acids of which 20 % are asparagines and contains an N-terminal signal peptide sequence (**Figure 2a**). Structural analysis using Alphafold ^34^ revealed that the protein was disordered with the exception of the signal peptide. In our scRNA-seq data, *STF1* was not only expressed in susceptible RHs pre-inoculation but also in other tissues like the cortex, meristem and pericycle (**Figure 1j, Supplemental Figure S3**). Moreover, its expression was further induced in RHs post rhizobia perception. We isolated three different *stf1 LORE1* alleles ^35^ to study its functional relevance. *stf1* plants displayed a wt-like onset of nodule organogenesis at 12 days post inoculation (dpi) with *M. loti* strain R7A (**Supplemental Figure S6**), and a hypernodulation phenotype with exclusively white nodules at 28 dpi (**Figure 2b, d**). This phenotype is commonly observed for mutants that fail to establish infected functional nodules like *rinrk1* ^36^, which prompted us to quantify the density of ITs. *stf1* plants displayed a 90 % reduction in IT density (**Figure 2d**). Many RHs had large infection pockets indicating successful initial perception of rhizobia but impaired IT progression (**Figure 2e**). We then tested nodulation phenotypes of *stf1* roots when inoculated with *Agrobacterium pusense* IRBG74, which enters Lotus roots via crack entry ^37^, or the more aggressive *M. loti* strain MAFF-dsRed (**Figure 2f, g**). Similar to R7A inoculation, *stf1* formed more white nodules than the wt independent of the rhizobia strain applied. However, all three *stf1* alleles also formed a few light and dark pink nodules with the more aggressive strains. In MAFF inoculated roots, ITs were able to penetrate *stf1* nodule primordia but bulged afterwards resulting in failed establishment of functional symbiosomes (**Figure 2h**). Aside from *STF1*, we also studied the function of the homologous gene *LotjaGi3g1v0447100* (*STF2*), which had a similar expression profile to *STF1* (**Supplemental Figure S7a**). *stf2* single mutants, however, displayed neither a nodulation nor an IT phenotype. *stf1,2* double mutants phenocopied *stf1* plants (**Supplemental Figure S7b**).

**Figure 2.**
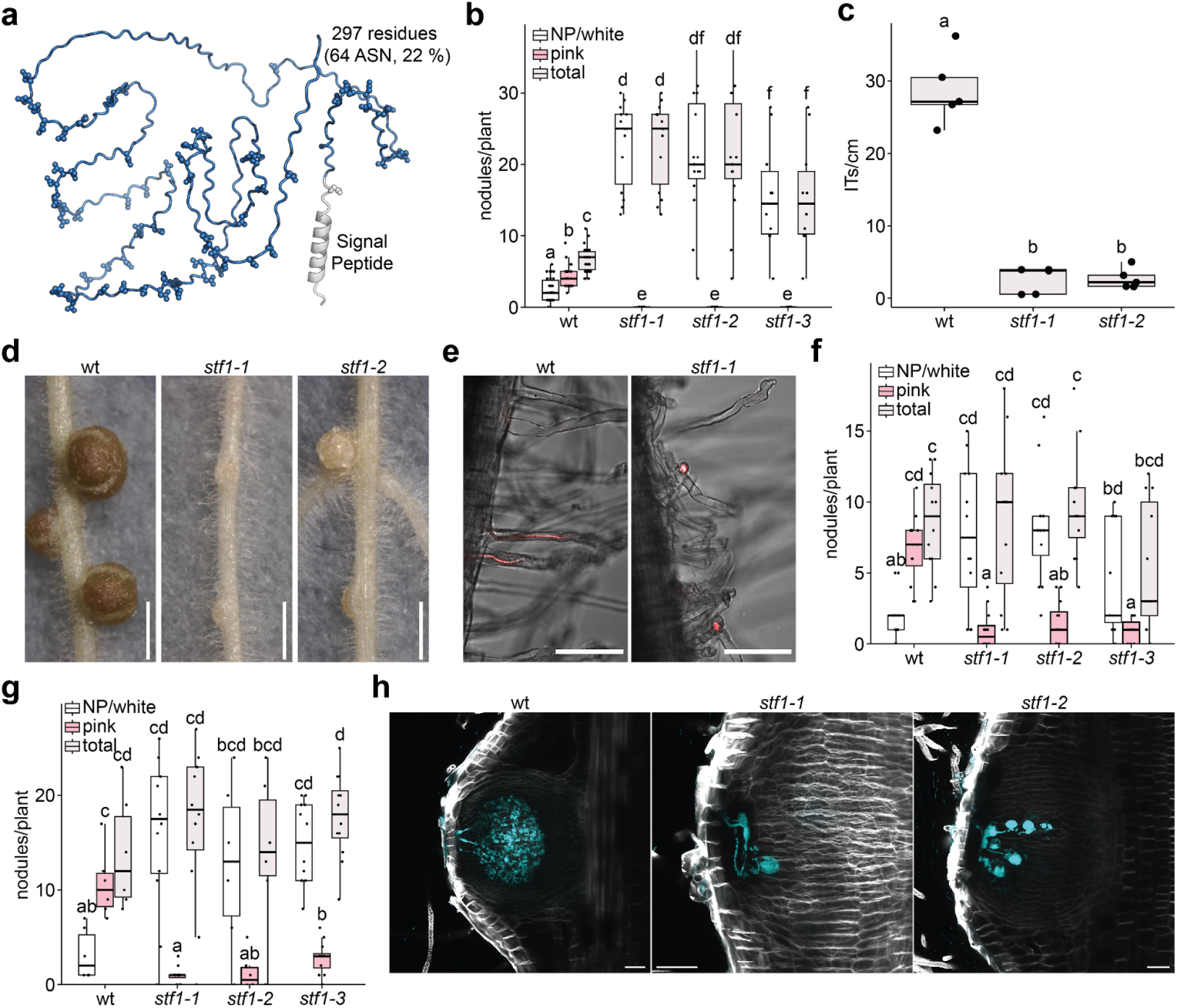
*STF1* is essential for normal IT formation. **a**) Predicted structure of STF1 with the N-terminal signal peptide and the 64 asparagines indicated. **b**) Number of white, pink and total nodules at wild-type (wt), *stf1-1, stf1-2* and *stf1-3* plants 28 days post inoculation (dpi) with *M. loti* R7A-dsRed (p ≤ 0.05; Scheirer–Ray–Hare with Benjamini-Hochberg-corrected Wilcoxon rank sum test; n ≥ 10 per genotype). **c**) IT density at wt, *stf1-1* and *stf1-2* plants 10 dpi with *M. loti* R7A-dsRed (p ≤ 0.05; one-way ANOVA; n = 5 per genotype). **d**) Pictures of representative wt, *stf1-1* and *stf1-2* plants 35 dpi with *M. loti* R7A-dsRed. Scale bars: 1 mm. **e**) Pictures of representative wt and *stf1-1* RHs with infection pockets or ITs 10 dpi. Scale bars: 100 µm. **f**) Number of white, pink and total nodules at wt, *stf1-1, stf1-2* and *stf1-3* plants 31 dpi with *A. pusense* IRBG74 (p ≤ 0.05; Scheirer–Ray–Hare with Benjamini-Hochberg-corrected Wilcoxon rank sum test; n = 12 per genotype). **g**) Number of white, pink and total nodules at wt, *stf1-1, stf1-2* and *stf1-3* plants 31 dpi with *M. loti* MAFF-dsRed (p ≤ 0.05; Scheirer–Ray–Hare with Benjamini-Hochberg-corrected Wilcoxon rank sum test; n ≥ 6 per genotype). Letters indicate significantly different statistical groups. **h**) Maximum intensity projections of representative wt, *stf1-1* and *stf1-2* nodules 8 dpi with *M. loti* MAFF-dsRed. Images show cell walls stained with Calofluor White (grey) and dsRed-labelled bacteria (cyan). Scale bars: 50 µm. All raw data and statistical analysis for the displayed experiments are available in Source Data “Supp data S3 - nodule and IT quantifications”.

### *NFR1* expression from susceptibility gene promoters is sufficient for root hair infection

Next, we asked how the susceptibility gene expression was related to expression of the *NPL* gene, which marks RH cells undergoing rhizobium infection ^28,29^. Using nucleus-localised mCHERRY driven by the previously published *NPL* promoter ^28^ combined with nucleus-localised tYFP driven by *EXPB1* or *STF1* promoters, we found that all *NPL::nls-mCHERRY* expressing cells also expressed *EXPB1::tYFP* or *STF1::tYFP*. No cells expressed *NPL::nls-mCHERRY* alone (**Figure 3a-c, Supplemental Figure S5**). Combined with the scRNA-seq data, these observations suggested that receptor-mediated NF perception restricted to susceptible cells should be sufficient to allow rhizobium infection. To test this prediction, we expressed *genomic NFR1* (*gNFR1*) under control of either its native promoter or the susceptible RH-specific promoters *EXPB1* or *STF1* in *nfr1* hairy roots. *EXPB1::gNFR1* rescued *nfr1* RH IT formation but not nodule formation. *STF1::gNFR1* rescued both IT formation and nodulation in the *nfr1* mutant (**Figure 3e, f**). The IT number in both cases was similar to that of wild-type hairy roots transformed with an empty vector. *nfr1* plants transformed with *NFR1::gNFR1* showed increased nodulation and infected thread counts compared to the wild type empty vector control, perhaps reflecting an incomplete *NFR1* promoter construct (**Figure 3f**). The *EXPB1* and *STF1* promoter rescue results are consistent with broader *STF1* expression in other cell types than RHs, compared to the much more restricted *EXPB1* expression (**Supplemental Figs. S3-5**), and confirm that driving *NFR* expression from susceptibility gene promoters is sufficient to allow rhizobium infection in RHs.

**Figure 3.**
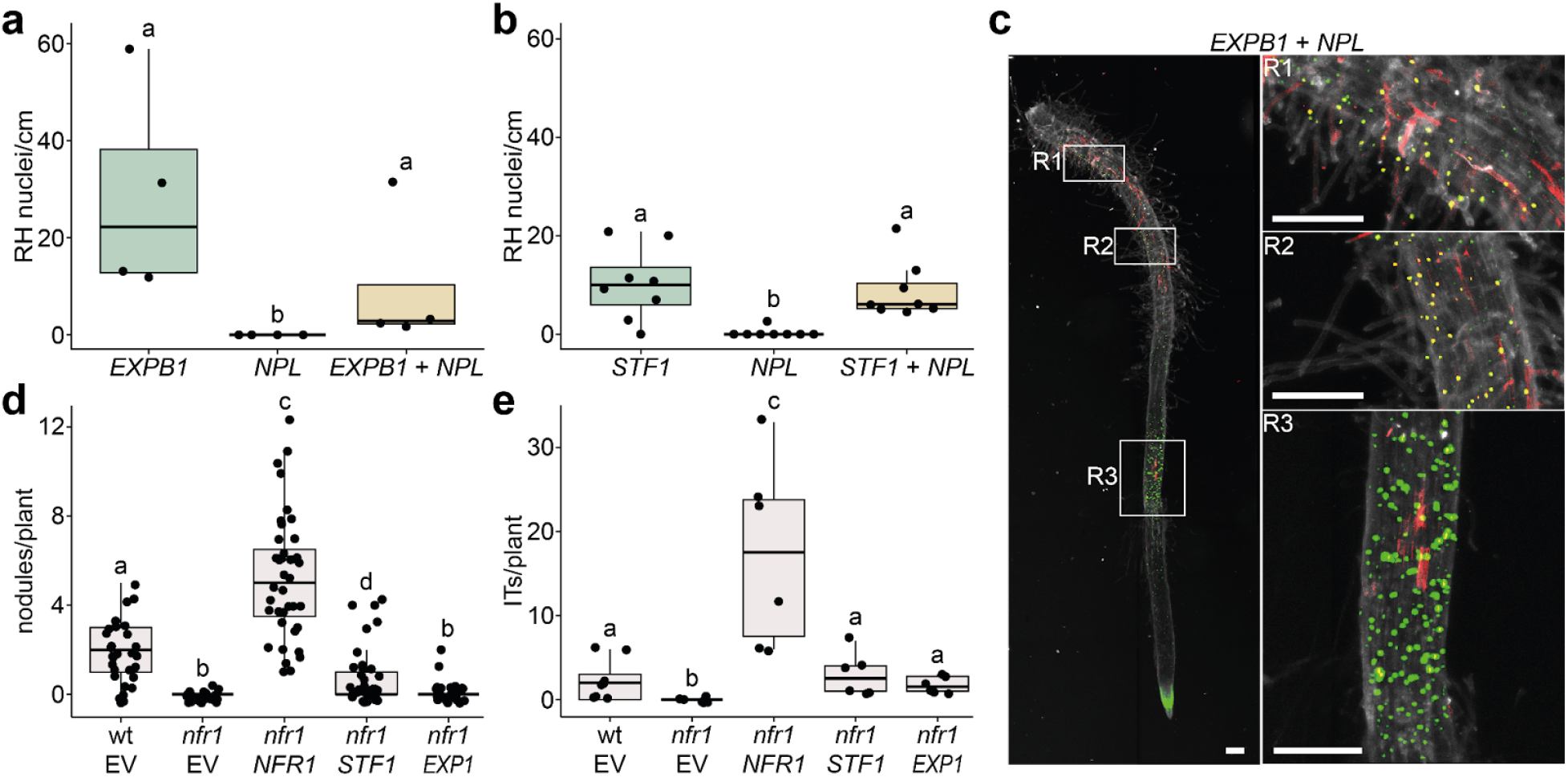
Susceptible RH marker gene expression coincides with *NPL* expression in RHs. **a**) Quantification of RH nuclei in hairy roots expressing *EXPB1::tYFPnls* and/or *NPL::nls-mCHERRY* 6 days post inoculation (dpi) (p ≤ 0.05; one-way ANOVA; n = 4 per group). **b**) Quantification of RH nuclei in hairy roots expressing *STF1::tYFPnls* and/or *NPL::nls-mCHERRY* 5 dpi (p ≤ 0.05; one-way ANOVA; n = 8 per group). **c**) *EXPB1* (tYFPnls reporter construct shown in green), *NPL* (nls-mCHERRY reporter construct shown in red) and *EXPB1-NPL* coexpression (shown in yellow) expression in wt hairy roots 5 dpi with *M. loti* R7A. White boxes indicate three magnified regions (R1-3) showing epidermal cells with exclusive *EXPB1* expression (R3, green) or overlapping *EXPB1* and *NPL* expression (R1 and R2, yellow). Scale bar: 100 µm. Single-channel images are depicted in Supplemental Figure S5. **d**) Nodule number of wild-type (wt) and *nfr1* hairy roots transformed with an empty vector (EV), or a vector containing *NFR1::gNFR1* (*NFR1*), *STF1::gNFR1* (*STF1*) or *EXPB1::gNFR1* (*EXPB1*) 35 dpi (p ≤ 0.05; Kruskal-Wallis followed by Benjamini-Hochberg-corrected pairwise Wilcoxon rank-sum test; n ≥ 23 per group). **e**) IT number of wt and *nfr1* hairy roots transformed with an empty vector (EV), or a vector containing *NFR1::gNFR1* (*NFR1*), *STF1::gNFR1* (*STF1*) or *EXPB1::gNFR1* (*EXPB1*) 21 dpi (p ≤ 0.05; Kruskal-Wallis followed by Benjamini-Hochberg-corrected pairwise Wilcoxon rank-sum test; n ≥ 6 per group). Letters indicate significantly different statistical groups. All raw data and statistical analysis for the displayed experiments are available in Source Data “Supp data S3 - nodule and IT quantifications” and “Supp data S4 - nuclei quantification”.

### Susceptible RH formation is conserved among legumes

After establishing the importance of susceptible RHs in Lotus, we studied the conservation of susceptible RHs among legumes in more detail. First, we performed a phylogenomic analysis of Lotus RH susceptibility marker gene orthologues within 42 non-nodulating and 45 nodulating species of the nitrogen-fixing nodulation (NFN) clade, and 40 other land plant species as an outgroup (**Supplemental Figure S8, Source Data “Supp data S5 - Phylogenomics”)**. We observed that RH susceptibility marker orthogroups appeared to be enriched in nodulating species compared to non-nodulating species (**Figure 4a, Source Data “Supp data S5 - Phylogenomics”**). The permutation test confirmed that this enrichment was significant compared to a random selection of RH marker genes (**Figure 4b**, p = 0.007, **Source Data “Supp data S5 - Phylogenomics”**). Next, we analysed a published scRNA-seq dataset from a distantly related legume species. We picked a recently published Medicago susceptible zone snRNA-seq dataset ^38^ (**Figure 4c**). We applied the same approach as for Lotus and clustered the 1033 RH cells into 18 subclusters (**Figure 4d**). Based on the gene expression of known NF-responsive genes, subcluster 0 was identified as infection-related (**Figure 4e, Source Data “Supp data S6 - Medicago RH gene lists”**). It contained 157 cells of which 32 belonged to control samples representing 19.5 % of the control RHs (n = 164 control RH cells, **Figure 4f**). The 19.5% susceptible RHs in the susceptible zone of Medicago root is substantially higher than the 0.7% we identified in Lotus whole roots, consistent with enrichment of ^38^. As for Lotus, susceptibility marker genes were highly specific (**Figure 4g**). We identified 392 susceptibility marker genes of which we could identify Lotus orthologues for 279. We used these 279 genes and the 163 Lotus susceptibility marker genes to test for a significant enrichment of marker genes in Lotus and Medicago susceptible RHs. The overlap of 9 genes displayed a significant enrichment (Fisher’s exact test p = 7.72e^-05^, **Source Data “Supp data S7 - Fishers exact test SRHs”**). The list of overlapping susceptibility marker genes comprised *MtrunA17Chr7g0231221* (*EXPB2*) which is known to determine IT formation in soybean ^39^, *MtrunA17Chr1g0203441* (*MYB74*), *MtrunA17Chr3g0128791, MtrunA17Chr2g0322651, MtrunA17Chr1g0202421, MtrunA17Chr8g0344521, MtrunA17Chr8g0393021, STF1* as well as *CBS1*. The latter was highly specific under control conditions and was induced further upon NF treatment in Medicago and R7A treatment in Lotus (**Figure 4h, i**). All in all, the two independent approaches indicate that susceptible RHs are conserved among legumes.

**Figure 4.**
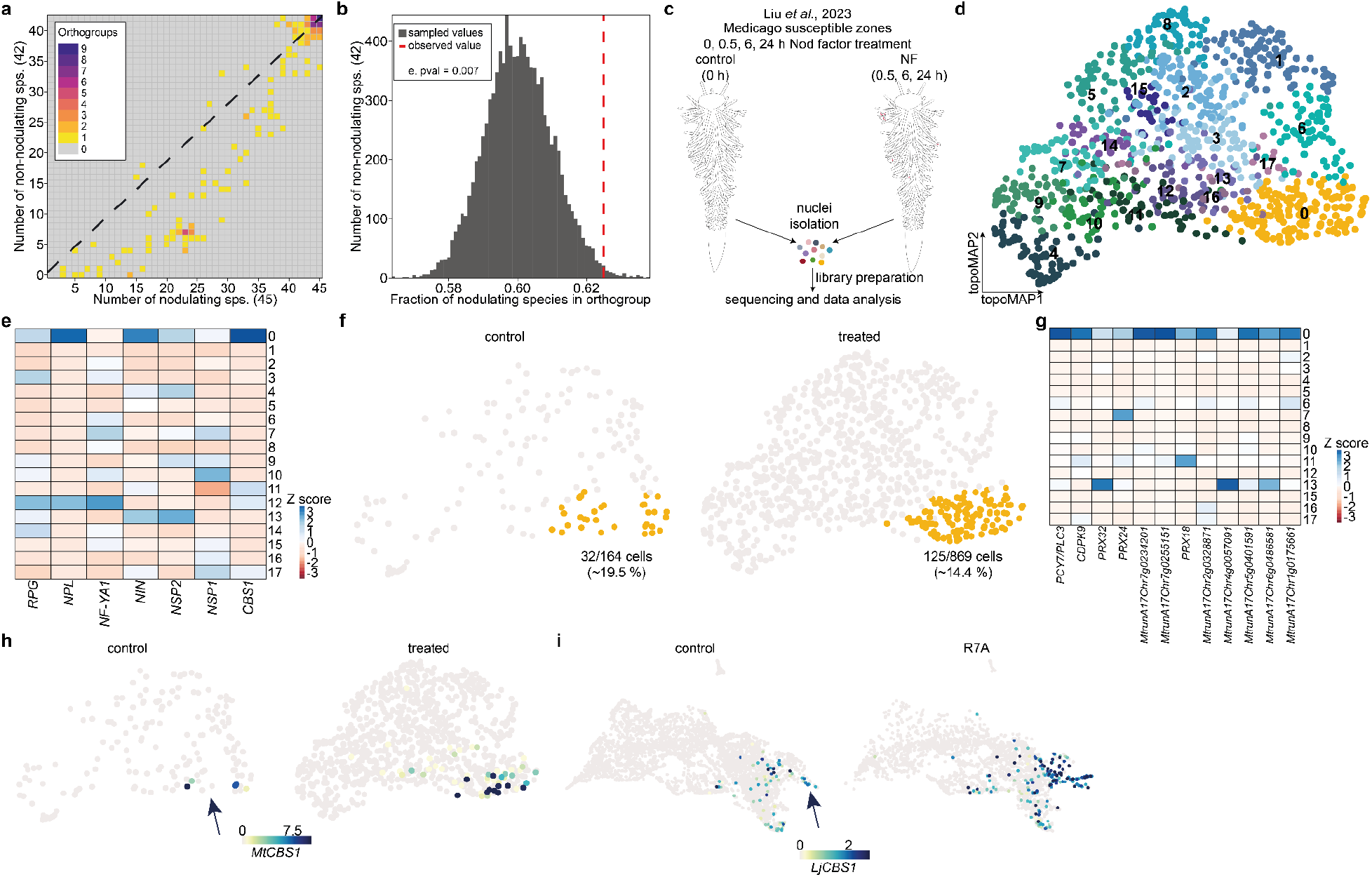
Susceptible RH formation is conserved in nodulating species. **a**) Occurrence of Lotus susceptibility gene (c21) orthogroups in nodulating (x-axis) and non-nodulating species (y-axis). The diagonal line indicates an 1:1 ratio between nodulating and non-nodulating species while the color gradient from yellow to purple depicts the number of orthogroups. **b**) Permutation plot for the occurrence of randomly picked RH marker gene sets (dark grey) against susceptibility genes (dotted red line) in nodulating and non-nodulating species. **c**) Summary of experimental procedure for Medicago snRNA-seq data generation in Liu *et al*., 2023 ^38^. *Medicago* susceptible zones were treated with Nod-factors (NF) and harvested 0, 0.5, 6 and 24 hours post treatment. 0 hpt are depicted as control while the other time points are summarised as “NF”. **d**) topoMAP of all Medicago RHs which were clustered into 18 subclusters. **e**) Expression of known infection-related genes in RH subclusters. **f**) RH cells split by treatment. Subcluster 0 containing susceptible and NF-responsive RHs is colored in gold while all other RH cells are depicted in grey. Percentage indicates the proportion of subcluster 0 cells compared to all RHs in one treatment. **g**) Expression patterns of the top12 subcluster 0 control marker genes in RH subclusters under control condition. **h**) Normalised Mt*CBS1* expression in Medicago RHs. **i**) Normalised Lj*CBS1* expression in Lotus RHs. All gene lists related to this data set can be found in Source data “Supp data S6 - Medicago RH gene lists”.

### The number of susceptible RHs correlates with the occurrence of infection events

After establishing the functional relevance and conservation of susceptible RHs, we speculated that the number of susceptible RHs might vary in symbiotic mutants and could be a predictor of the number of successful infection events. Therefore, we generated the *har1 ein2a,b* triple mutant as a Lotus analogue of the Medicago *skl sunn* double mutant ^40^. Similar to *sklsunn, har1 ein2a,b* hypernodulated and displayed a 3.5-fold increased IT density compared to wt (**Figure 5a, Supplemental Figure S9**). We then applied scRNA-seq to see how many susceptible RHs we could identify and integrated all *har1 ein2a,b* cells with the wt integrated object and extracted the RH cells. Compared to the number of susceptible RHs in wt roots, 3-times more susceptible RHs were annotated in *har1 ein2a,b* under control conditions correlating with the 3.5-fold increase in IT density (**Figure 5b**). Compared to wt, more *har1 ein2a,b* susceptible RHs expressed *EXPB1* and the expression was stronger in those susceptible RHs (**Figure 5c**). To dissect whether autoregulation of nodulation or ethylene signalling are causative for the increase in susceptible RHs and to confirm the scRNA-seq data, we quantified the number of susceptible RHs by using the *EXPB1* reporter in wt, *har1, ein2a,b* and *har1 ein2a,b* hairy roots (**Figure 5d**). Compared to wt, over 1.8-times more *ein2a,b* and *har1 ein2a,b* RHs expressed *EXPB1*, providing *in planta* confirmation for the observations in the scRNA-seq analysis and IT quantification and demonstrating that ethylene signalling is a major regulator of susceptible RH formation. Consistent with these observations, genotype density means for EXPB1-positive RHs and IT density showed a strong positive correlation (Spearman ρ = 1.0 across four genotypes; **Source Data “Supp data S8 - Spearman”**), with both measures increasing from wild type through *har1* and *ein2a,b* to *har1 ein2a,b* (**Supplemental Figure S10**).

**Figure 5.**
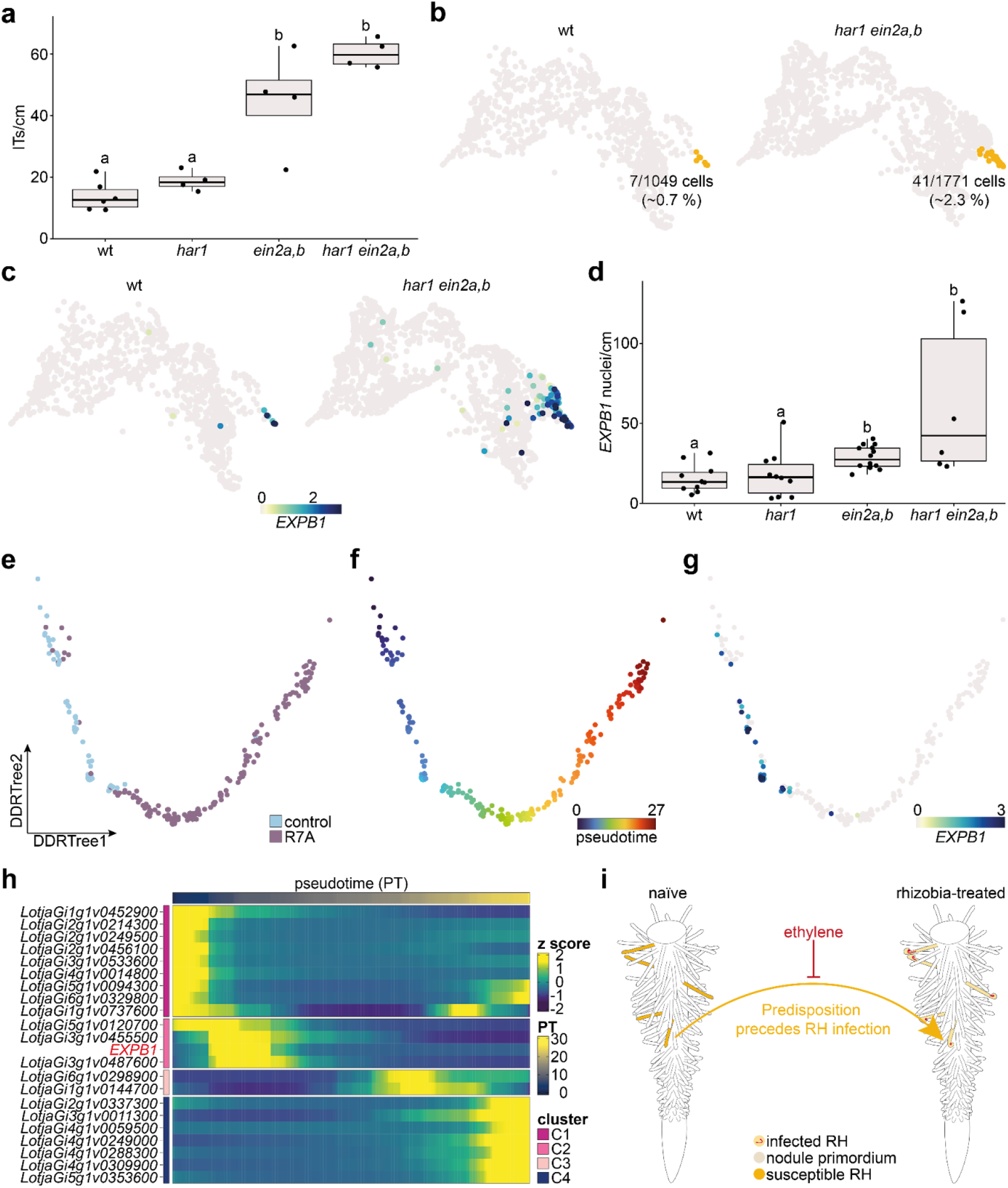
The number of susceptible RHs is correlated with infection density in symbiotic mutants. **a**) IT density at wt, *har1, ein2a,b* and *har1 ein2a,b* plants 8 dpi with *M. loti* R7A-dsRed (p ≤ 0.05; one-way ANOVA; n ≥ 4 per genotype) **b**) TopoMAP of control wt and *har1 ein2a,b* RHs 1 dpt with water. Gold: cell cluster containing susceptible RHs and rhizobia-responsive RHs; Grey: all other RH cells. **c**) Normalised *EXPB1* expression in scRNA-seq. **d**) Quantification of *EXPB1::tYFPnls*-expressing RH nuclei in wt, *har1, ein2a,b* and *har1 ein2a,b* control hairy roots (p ≤ 0.05; Kruskal-Wallis followed by Benjamini-Hochberg-corrected pairwise Wilcoxon rank-sum test; n ≥ 6 per genotype). Letters indicate significantly different statistical groups. **e**,**f**) DDRTree map of all wt and *har1 ein2a,b* cluster 21 cells colored by **e**) treatment, and **f**) Monocle2 pseudotime. **g**) Normalised *EXPB1* expression in cluster 21 RHs and **h**) Gene expression of trajectory cluster markers over Monocle2 pseudotime. **i**) Model of ethylene-dependent susceptible RH establishment as a fundament for RH infection. RH susceptibility could be regulated by other phytohormones, microbes, nutrients or soil properties. All raw data and statistical analysis for the displayed experiments are available in Source Data “Supp data S3 - nodule and IT quantifications” and “Supp data S4 - nuclei quantification”.

Ultimately, we performed pseudotime analysis to study the trajectory of RH infection. In a pseudotime analysis, cells are ordered along a pseudotime axis based on their transcriptome, simulating temporal progression. We hypothesised that if susceptible RHs are the initial state of successfully infected RHs, they should cluster at an early point in pseudotime, while further progressed infected RHs should cluster at later pseudotimes. We first attempted to perform this analysis with only wt RH cells but did not succeed so due to the low number of susceptible RHs. We then combined all wt and *har1 ein2a,b* cells and performed the pseudotime analysis with Monocle 2 ^41^. In the resulting DDRTree maps, cells were separated by treatment and control cells were found at early pseudotimes (**Figure 5e, f**). *EXPB1* was expressed early in the trajectory and disappeared with pseudotime progression (**Figure 5g, Supplemental Figure S11**). In total, four clusters were identified along the trajectory, the first of which appeared to be unspecific, associated with genes broadly expressed in many RHs (**Figure 5h, Supplemental Figure S11a-b**). The second cluster displayed early susceptibility and included *EXPB1* as a marker gene. Moreover, clusters 3 and 4 represented later infection time points (**Supplemental Figure S11**). Taken together, our hormone mutant and pseudotime analysis showed that RH susceptibility is negatively regulated by the gaseous hormone ethylene and supported that susceptible RHs represent the initial state of successfully infected RHs (**Figure 5i**).

## Discussion

In this study, we identify a rare, transcriptionally distinct population of RHs that are pre-specified for rhizobial infection before bacterial contact. These susceptible RHs, which represent only a small fraction of all RHs, express infection-related genes prior to NF exposure and induce the expression of essential infection genes further to accommodate rhizobia after NF perception. This finding challenges the prevailing view that infection competence arises *de novo* in response to bacterial signals and instead reveals that legumes pre-allocate infection capacity to a dedicated cell population.

The expression of canonical infection markers including *CBS1* ^24^, *EXPB2* ^39^, *EXPA1* ^26^ and the novel gene *STF1* in susceptible RHs in the absence of rhizobia (**Figure 1**) demonstrates that legumes restrict symbiotic engagement through cell state specification rather than through uniform receptor distribution and purely signal-driven activation. Although *NFRs* are broadly expressed across the RH population ^3,4^, only susceptible RHs progress to infection following bacterial perception. The observation that all rhizobia-responsive cells derive from this pre-existing susceptible population (**Figure 3**) provides direct evidence that predisposition, rather than rhizobial induction alone, determines which cells become infected. Similar patterning principles have been proposed in arbuscular mycorrhizal symbiosis, where cortical cell competence for fungal accommodation is established before colonization, but after fungal contact ^42,43^, and in the legume cortex, where chromatin remodelling at symbiosis genes creates a permissive state for nodule organogenesis after rhizobia perception ^44–46^.

The functional requirement for *STF1*, encoding a disordered E6-like protein with an N-terminal signal peptide, reveals unexpected molecular players in IT formation within the susceptible RHs. The strong reduction in IT density, accumulation of infection pockets and impairment of infection events via crack entry in *stf1* mutants (**Figure 2**) indicate that STF1 acts upstream of sustained infection progression in RHs, epidermis and cortex. Together with the known roles of STF1-like proteins in regulating cell wall dynamics during fiber elongation, pollen-stigma interaction and fruit ripening ^31–33^, these findings support a model in which STF1 could modify extracellular matrix properties of susceptible RHs, epidermal cells and the cortex to facilitate bacterial entry and IT elongation. The intrinsically disordered nature of STF1 could indicate a flexible scaffold or spacer function ^47^ that could reorganize cell wall architecture at the infection site, although direct biochemical and biophysical characterization will be required to test this model.

The quantitative relationship between susceptible RH abundance and infection density across genotypes provides a framework for understanding how host signalling pathways tune symbiotic capacity. The hyperinfection phenotypes of *ein2a,b* double and *har1 ein2a,b* triple mutants correlate with an increased frequency of susceptible RHs (**Figure 5, Supplemental Figure S10**), demonstrating that ethylene signalling shapes infection capacity by regulating susceptible RH specification rather than acting solely on post-perception responses. In this view, ethylene integrates multiple environmental inputs such as pH, hypoxia, and nutrient availability ^48–50^, raising the possibility that susceptible RH formation adjusts dynamically to soil conditions. Whether susceptible RHs represent a stable developmental end point or a transient physiological state remains unresolved.

The conservation of susceptible RH populations among nodulating species (**Figure 4**), including shared expression of *STF1* and *CBS1* in Lotus and Medicago, suggests that this mechanism arose early in legume evolution and is under strong selective pressure. Spatially restricting infection to a minority of RHs may serve as a safety mechanism to limit potential exploitation by non-beneficial microbes. The induction of susceptibility-associated markers like *RHD6LA* in response to non-symbiotic rhizobacteria ^22^ raises the possibility that susceptible RHs form part of a more general microbial surveillance architecture, with subsequent signalling modules - including calcium signalling dynamics and cortex-specific chromatin remodelling - determining whether interactions resolve toward symbiosis or defence.

Methodologically, the identification of susceptible RHs, and a specific set of associated genes, provide powerful tools for dissecting symbiotic signalling with high spatial and temporal resolution. The sufficiency of *EXPB1* and *STF1* promoters to drive *NFR1* expression and rescue infection in *nfr1* mutants demonstrates that manipulating a small pre-specified cell population is sufficient to restore infection capacity. This cell-type-specific engineering strategy complements existing approaches that broadly modulate signalling pathways and offers a route to transfer infection competence to non-legume species by reconstructing a defined susceptible cell state. Our work also raises new questions for future study. First, the upstream regulators that specify susceptible RHs remain unknown. Analysing an enrichment of specific transcription factor motifs in susceptibility marker promoters might point to candidate regulators, awaiting subsequent genetic and biochemical validation. Second, this study primarily characterizes transcriptional signatures. Additional layers such as chromatin accessibility, epigenetic regulation, and metabolic rewiring could further define the susceptible state, paralleling how PRIMER cells and symbiotic cortical cells are distinguished by combined transcriptomic and chromatin features ^44,51^. Third, environmental plasticity of susceptible RH formation is not yet resolved: systematic variation of nitrogen availability, microbial community composition, and abiotic stress will be required to determine how flexible this state is and whether it can be reprogrammed in agronomic contexts. Finally, while the conservation in Lotus and Medicago is clear, it remains unknown whether analogous rare cells, developmentally primed for symbiosis with actinorhizal or mycorrhizal fungi exist in non-legumes.

Together, our findings show that legumes proactively allocate symbiotic competence to a rare RH population that initiates and sustains rhizobial infection. Our work supports the emerging paradigm in which rare, spatially confined cells act as gatekeepers and organizers of plant-microbe interactions. At a practical level, understanding the transcriptional makeup of specific cells pre-disposed for symbiont accommodation, offers new opportunities for engineering analogous symbiont accommodation in non-legume crops.

## Methods

### Plant material

For *M. loti* R7A nodulation assays, IT density quantification, and scRNA-seq, seeds of *Lotus japonicus* accession Gifu and respective mutants were scarified with sandpaper, sterilised in an 1 % (v/v) NaClO solution for 12 minutes, washed three times with sterile ddH_2_O and germinated on filter paper under long-day conditions (8h dark/16h light) with a light intensity of 120 µmol m^−2^ s^−1^ at 21°C. Three days after germination, seedlings were transferred to square plates with 1.4% Agar Noble slopes containing 0.25x B&D medium covered with filter paper. A metal bar with 3-mm holes for roots was inserted at the top of the agar slope to shade roots from light during cultivation. For nodulation assays and IT density quantification, seedlings were inoculated with 500 µL of either water (control) or *M. loti* R7A (R7A, OD_600_ = 0.02) along the length of the root. All plants were cultivated under long-day conditions (8h dark/16h light) with a light intensity of 120 µmol m^−2^ s^−1^ at 21°C.

For nodulation assays and microscopy with *M. loti* MAFF-dsRed and nodulation assays *A. pusense* IRBG74-dsRed, seeds were scarified with sand paper and sterilised for 3 minutes using 3% (v/v) NaClO, followed by several washes in sterile dH_2_O and imbibition in sterile dH_2_O for 3 days at 10°C, agitating. Three days post germination on 1% water agar, seedlings were transferred to 0.25x B&D agar with filter paper. Seeds were germinated and plants grown at long-day conditions (8h dark/16h light), with a light intensity of 120 µmol m^−2^ s^−1^ at 21°C, and with roots shaded from light. Inoculation with *M. loti* MAFF-dsRed, or IRBG74-dsRed was performed 7 days post germination with a bacterial OD_600_ of 0.02, using 75 µL of bacterial suspension per root. For genetic studies, *nfr1-1* ^4,52^, *har1* ^53^, *ein2a,b* ^54^, *har1 ein2a,b*, as well as LORE lines *stf1-1* (30090214), *stf1-2* (30090226), *stf1-3* (30164854) and *stf2-1* (30005945) were used ^35^. Primer sequences for *stf* genotyping can be found in Source Data “Supp data S9 - constructs primer promoter”.

### Rhizobia strains

To induce nodulation and IT formation, *L. japonicus* was inoculated with *M. loti* R7A (wt/dsRed/eGFP), which was cultured for 2 days at 28 °C on yeast mannitol broth (YMB) medium and resuspended to an optical density_600_ of 0.02 in sterile water. Similarly, inoculation with *M. loti* MAFF-dsRed and *A. pusense* IRBG74-dsRed was performed after culturing the bacteria for 2 days at 28°C in Yeast Extract Tryptone (TY) medium (supplemented with 1mM CaCl_2_) and resuspension of the bacteria to an OD_600_ of 0.02 in Farhäeus medium (supplemented with 1mM CaCl_2_).

### Protoplast isolation and single-cell RNA-sequencing

Protoplast isolation and scRNA-seq library preparation (Chromium Next GEM Single Cell 3’ Kit v3.1) were performed as described previously ^21^ using wt susceptible zones 5 dpt and wt and *har1 ein2a,b* whole roots (1 dpt) treated with water or inoculated with *M*.*loti* R7A as starting material.

### Raw data pre-processing, integration, clustering and pseudotime analysis

Recently published Gifu wt whole root 10 dpt and susceptible zone 5 dpt scRNA-seq raw sequencing data ^21,22^ and newly generated raw sequencing data were processed using Cell Ranger v6.1.2 (10X Genomics). The Lotus japonicus Gifu v1.2 and Gifu v1.3 genome assembly and gene annotations, respectively (available in Lotus Base, lotus.au.dk ^27^) were used as reference for “cellranger mkref”. “cellranger count” was run with the default parameters using the STAR 2.7.2a ^55^ aligner. The “filtered_feature_bc_matrix” was used as input for the analyses. The downstream analyses were carried out using Seurat 5.1.0 ^56^.

The gene count matrices were filtered to exclude any cells that had fewer than 200 or more than 7500 expressed genes and less than 500 UMIs. The cells were also filtered on the mitochondrial and chloroplast encoded gene expression, retaining only the cells expressing under 5% read counts from these features. Additionally, only genes that were expressed in at least three cells were included in the analysis.

All samples were normalised using the “SCTransform” function implemented in Seurat, with “vars.to.regress” set to mitochondrial and chloroplast genes. The samples were integrated using the canonical correlation analysis integration pipeline from Seurat, using the 10 dpt control samples as reference and selecting 3000 integration anchors.

PCA dimensionality reduction was run using the default function in Seurat on the integrated data assay. The functions “FindNeighbors” and “RunUMAP” were run using 50 principal components. The cells were clustered using the unsupervised Louvain clustering algorithm with a resolution of 1. The cell type for each cell was predicted based on the published Gifu wt whole root 10 dpi dataset. Clusters 4 and 18 were selected as the RH clusters based on the predicted annotations.

The TopOMetry ^23^ analysis for the whole dataset and RHs subset was performed on the scaled expressions for the 3000 genes in the integrated assay of the Seurat object. The analysis was run using the default command with the kernel set to ‘bw_adaptive’. The eigen methods used were ‘msDM’, ‘DM’ and the ‘MAP’ projection. The clustering resolution was set to 2.

Marker genes for TopOMetry clusters were identified using the FindMarkers function together with Presto ^57^, with p-value adjusted ≤ 0.05 (Lotus) and p-value ≤ 0.05 (Medicago).

Monocle2 ^41^ pseudotime trajectories were estimated for the lotus RH subcluster 21 using the “RunMonocle2” function from the scop ^58^ R package. The start point for the pseudotime ordering was manually selected.

### Gene Ontology analysis

Gene Ontology of susceptible RH markers was performed with topGO ^59^ using the Gifu 1.2 gene ontologies from Lotus Base ^27^.

### Promoter reporter constructs

Entry vectors containing *STF1* (2kb) and *EXPB1* (1.6 kb) promoter fragments flanked with BsaI sites were synthesised by Thermo Fisher and subsequently used for Golden Gate cloning of lvl and lvl2 *promoter:tYFPnls/nls-mCHERRY* and *nfr1* rescuing constructs. All construct names and promoter sequences can be found in Source Data “Supp data S9 - constructs primer promoter”

### Hairy root transformation

pIV101 vectors harboring *promoter:tYFPnls/nls-mCHERRY* or *nfr1* rescuing constructs were brought into Agrobacterium rhizogenes AR1193 as described previously ^60^. Hairy root formation of wild-type, *nfr1, har1, ein2a,b* and *har1 ein2a,b* plants was induced by piercing Lotus seedlings at the hypocotyl site by using a narrow needle with a drop of agrobacterium in ddH_2_O. Three weeks after inoculation with Agrobacterium the primary root of infected seedlings was cut, and plants exhibiting transgenic hairy roots were placed into lightweight expanded clay aggregate (LECA) containing 50 mL 1/4 B&D solution and were inoculated with *M. loti* (wt/eGFP) to promote nodulation and IT formation.

### Confocal microscopy

Confocal microscopy of hairy roots and *stf* mutant phenotyping using *M. loti* R7A wt/dsRed/eGFP was performed with a Zeiss LSM780 microscope. Hairy roots were either used directly for microscopy or submerged in (0.1% (w/v) Calcofluor White (CFW; Fluorescent Brightener 28, MP Biomedicals) for 1 minute and washed three times with water. The following excitation/emission [nm] settings were used: (i) Calcofluor White/autofluorescence of cell wall: 405/420-505, (ii) DsRed and mCHERRY: 561/580-660, (iii) tYFP: 514/517-560 (iv) eGFP: 488/500-520.

For *stf* mutant phenotyping using *M. loti* MAFF-dsRed, roots were fixed in 4 % (w/v) PFA (in PBS), for 2 × 5 min in vacuum and 45 min at room temperature (RT). After 3 washes of 5 min each in PBS, roots were stored in Clearsee, which was refreshed every 2 - 3 days. After CFW staining (0.1% (w/v) CFW in Clearsee) ^61^ for 45 min, roots were washed with and kept in Clearsee until imaging. All steps were performed at RT, roots were protected from light and gently agitated. Imaging was performed with a TCS SP8 confocal microscope (Leica Microsystems) controlled by the LAS X v3 software with λ_ex_= 405, λ_em_=425-475 for CFW detection and λ_ex_=561, λ_em_=570-599 for detection of dsRed-labelled bacteria. All images were processed using Fiji/ImageJ ^62,63^ and are displayed as overlaps of maximum intensity projections of the CFW and dsRed channel.

### Counting of fluorescent nuclei-containing RHs

To count nuclei of RHs, transgenic hairy roots expressing reporter constructs were analysed with a Zeiss LSM780 microscope. Only RHs at the sides of the hairy roots were counted and normalised against the respective hairy root length.

### Correlation analysis

To assess the association between susceptible RH abundance and infection capacity across genotypes, Spearman’s rank correlation coefficient between genotype means of EXPB1::tYFPnls-positive RH density and IT density was performed on four genotypes (wt, *har1, ein2a,b* and *har1 ein2a,b*) using the base cor.test function in R (version 4.5.1) with method set to “spearman”. Detailed values are provided in Source Data “Supp data S8 - Spearman”.

### Structural analysis of STF1 protein

The protein structure of Lotus STF1 was predicted using AlphaFold3 with default parameters ^64^. Five independent models were generated and the top-ranked model with the highest predicted confidence scores (pLDDT) was selected for structural analysis. Structural visualization was performed using PyMOL (The PyMOL molecular graphics system, version 2.5.2, Schrödinger, LLC).

### Phylogenomic analysis

To determine the presence of orthologs of genes related to RH susceptibility in members of the NFN clade, we gathered proteomes of 42 non-nodulating and 45 nodulating species, plus 40 other land plant species as outgroup (Source Data “Supp data S5 - Phylogenomics” sheet “genomes”). Orthologous groups were reconstructed using OrthoFinder v2.5.5 ^65^ with the parameter ‘-S diamond_ultra_sens’. An adjusted species tree was prepared (Supplemental Figure S8) based on the Angiosperm Phylogeny Website (v14) and relevant literature on relationships between different sub-linages of the NFN clade ^66–69^. This species tree was used for a second Orthofinder run using the parameters ‘-M msa’ and ‘-b’ to continue from previous blast results. Hierarchical orthologous groups (HOGs) were extracted for two nodes: (1) the common ancestor of the NFN clade and (2) the common ancestor of Fabales. The presence/absence of any gene in these HOGs was determined for all species using R v4.2.2 ^70^ and the tidyverse metapackage v2.0.0 ^71^. Any HOG that includes a *L. japonicus* marker gene expressed for the susceptible RH cell cluster (21) were selected, and the presence/absence of genes in those HOGs was plotted using the R package ggplot2 v3.5.2 ^72^. To determine whether the enrichment of these HOGs in nodulating species was due to chance, we made use of a random-sampling approach. Of the 163 genes significantly expressed, 158 were assigned to a NFN HOG. Thus, we randomly sampled 158 genes of the 4429 genes that are assigned to HOGs, and that were significantly expressed in any root-hair cluster, using the srswor (simple random sampling without replacement) function from the R package sampling v2.10 ^73^. For a total of 10,000 random samples, the NFN/Fabales HOGs of the sampled genes were extracted, and the mean fraction of nodulating species over the total number of NFN species with known nodulating status was calculated. The observed value was compared to distribution of these sampled values, and the empirical p-value was estimated as the proportion of random samples that showed a higher mean fraction of nodulating species than the observed number. Additionally, we did the same calculations with a “filtered” set, where before calculation, HOGs were excluded that appeared in five or fewer species, or that appeared in all species, as these are highly common and may bias the result. The same approach was done for all other root-hair clusters (Source Data “Supp data S5 - Phylogenomics” sheet “all_permutation_results”).

## Data availability

The data that support the findings of this study are available in the Supporting Information of this article. Sequencing raw data is deposited at the European Nucleotide Archive (ENA) with the Accession ID PRJEB108942.

## Acknowledgements

This work was supported by the European Research Council (ERC) under the European Union’s Horizon 2020 research and innovation programme (grant agreement no. 834221) and by the project Enabling Nutrient Symbioses in Agriculture (ENSA), that is funded by Gates Agricultural Innovations (INV-57461). The authors thank Beatrice Lace for her help/guidance during image acquisition of the *stf* mutant nodules.

## Author contributions

MF, HL, JSa, NA, MN, EBKS, FvB, HR and DR performed experiments. MF, LIF and ML performed scRNA-seq data processing. MF, LIF, ML, JSa, FvB, PMD, TO and SUA analysed and interpreted data. KRA performed structural analysis. MF and SUA conceptualised the research and wrote the first version of the manuscript. MF and SUA edited the manuscript with input from all authors.

## Supplemental Material

**Supplemental Figure S1.**
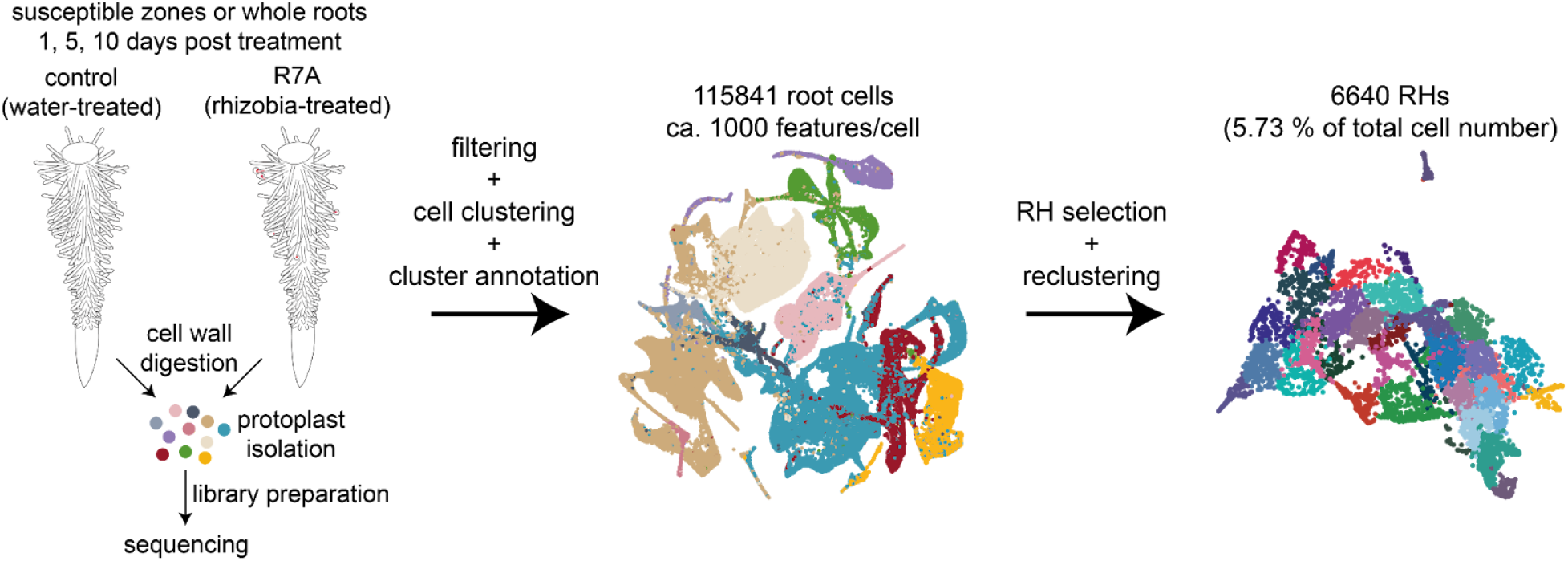
Overview over scRNA-seq data analysis. From left to right: Protoplasts of root and susceptible zone material were isolated and used for library preparation. After sequencing, raw sequences and low quality cells were filtered, 115841 high-quality root cells clustered based on their transcriptome and the resulting cell cluster annotated to a tissue type. RH cells were selected, reclustered and used to study susceptible RHs.

**Supplemental Figure S2.**
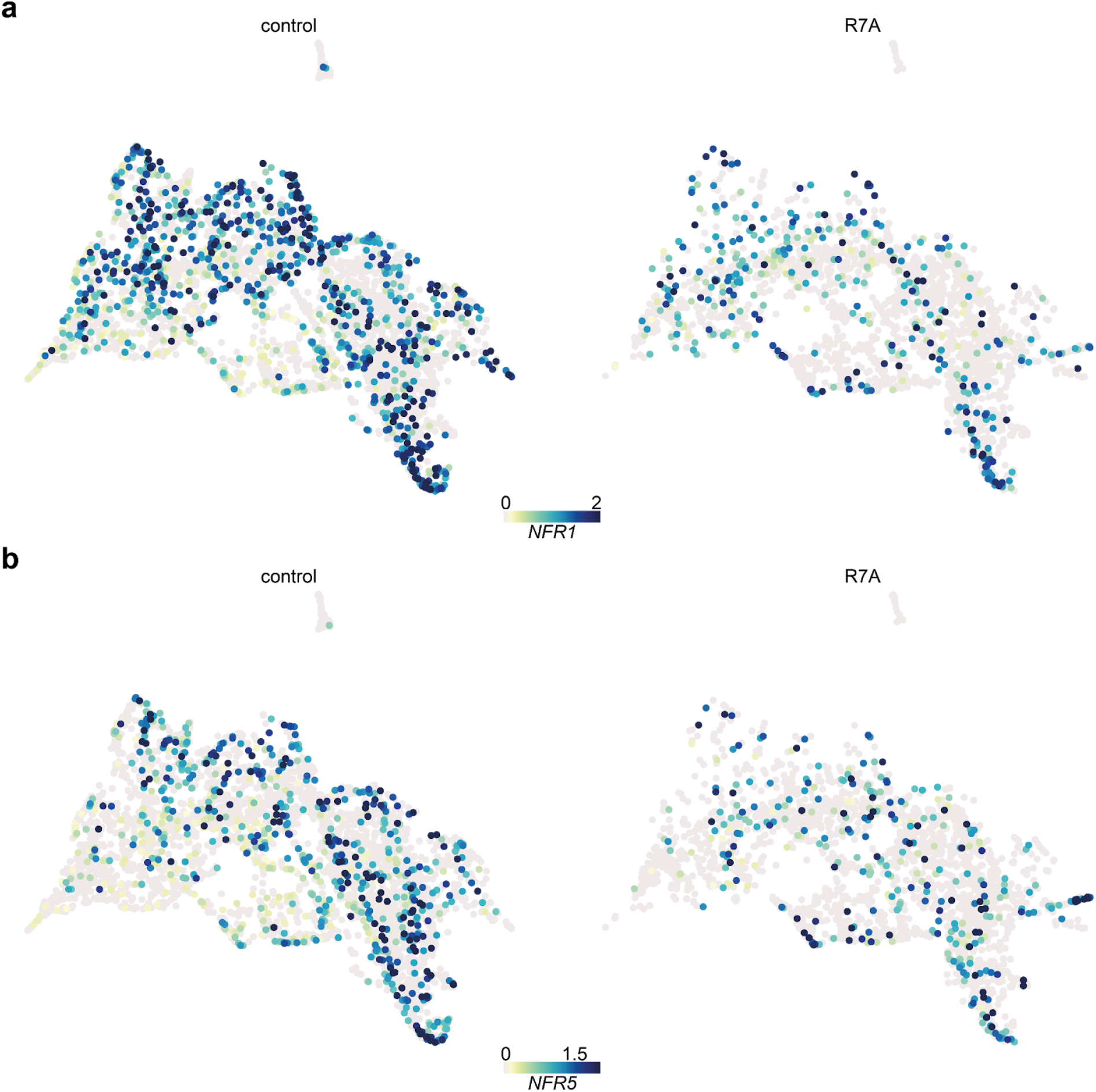
*NFR* expression in scRNA-seq. **a**) Normalised *NFR1* expression in all RH cells. **b**) Normalised *NFR5* expression in all RH cells.

**Supplemental Figure S3.**
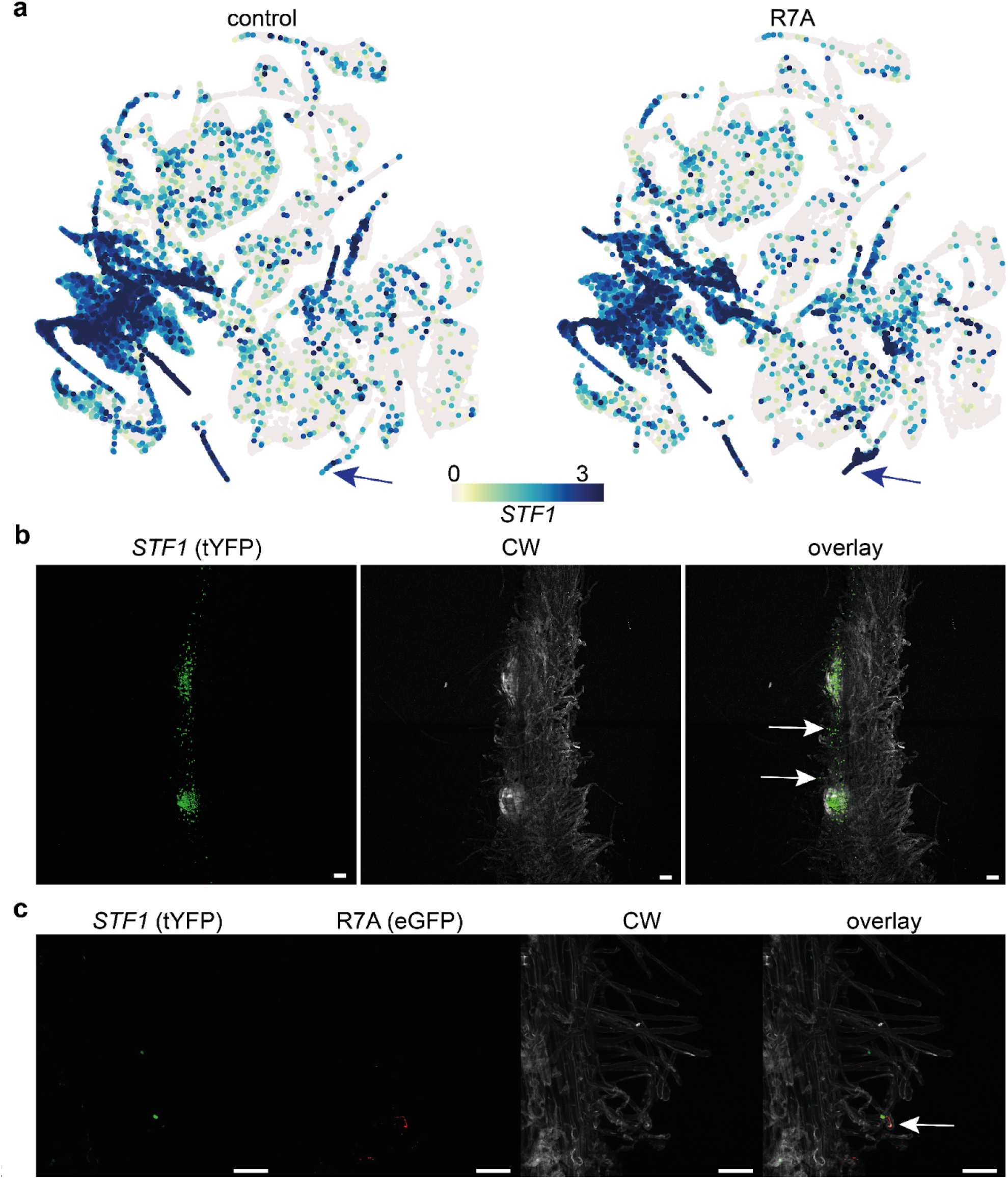
*STF1* expression in scRNA-seq and transgenic hairy roots. **a**) Normalised *STF1* expression in all wt cells. Arrows indicate responsive RHs. **b**) *STF1* expression (tYFPnls reporter construct shown in green) in wt control hairy roots. Arrows indicate RHs with fluorescent nuclei. Scale bar: 100 µm. **c**) *STF1* expression (tYFPnls reporter construct shown in green) in hairy roots inoculated with R7A-eGFP (shown in red) 7dpi. RH with IT and fluorescent nucleus is marked with an arrow. Scale bar: 100 µm.

**Supplemental Figure S4.**
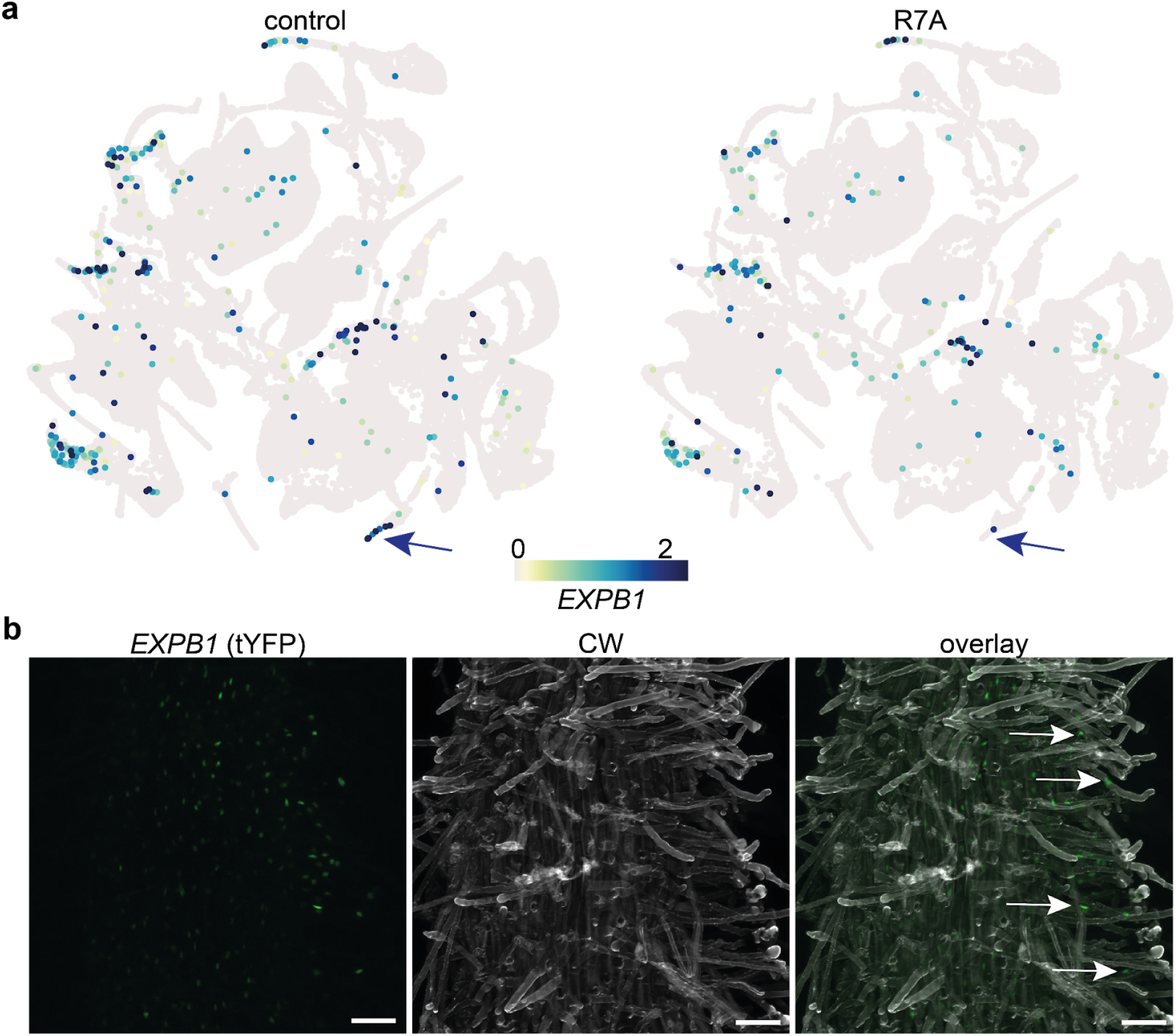
*EXPB1* expression in scRNA-seq and transgenic hairy roots. **a**) Normalised *EXPB1* expression in all wt cells. Arrows indicate rhizobium responsive RHs. **b** *EXPB1* expression (tYFPnls reporter construct shown in green) in wt control hairy roots. Arrows indicate RHs with green fluorescent nuclei. Scale bar: 100 µm.

**Supplemental Figure S5.**
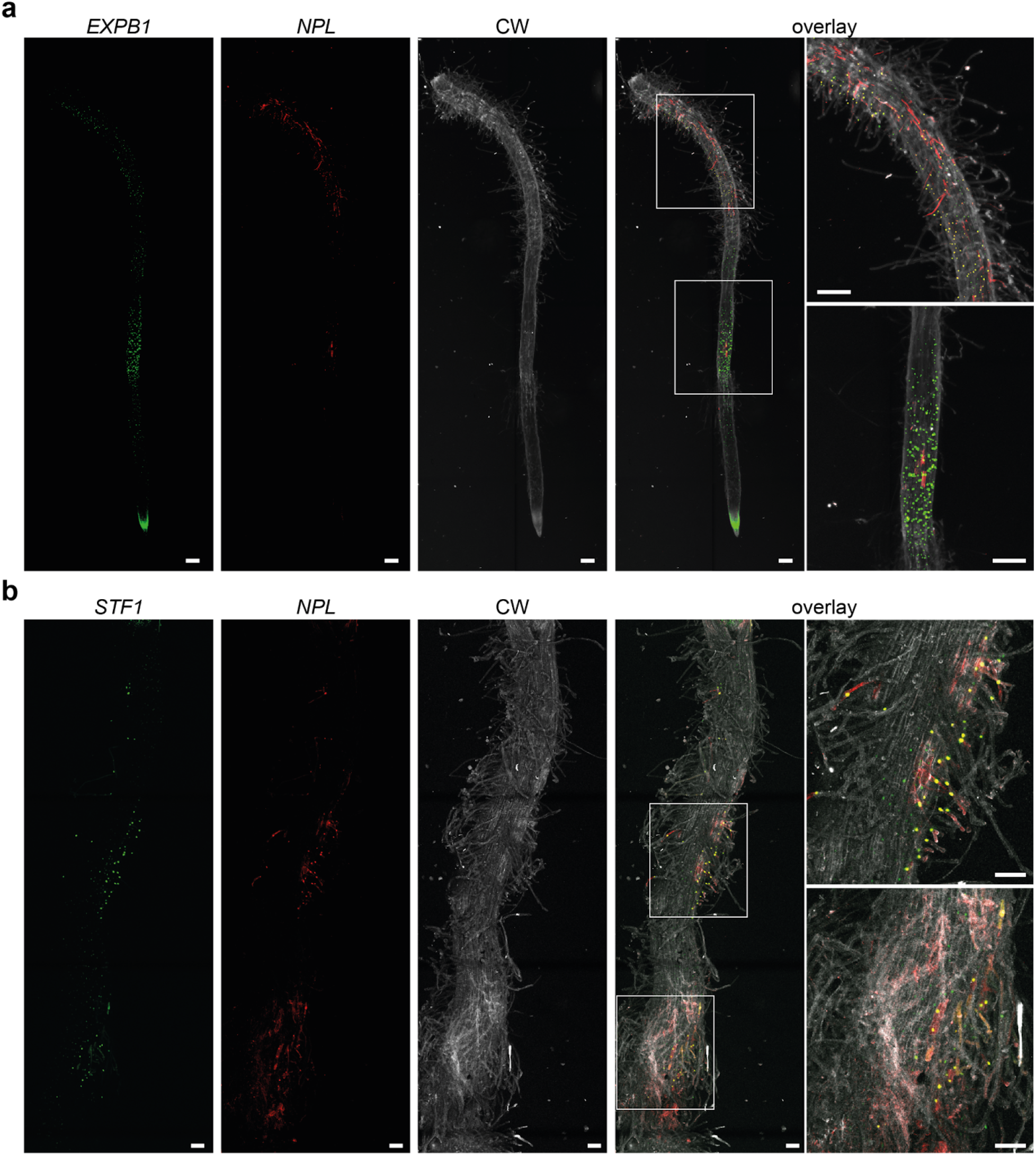
*EXPB1* and *STF1* coexpression with *NPL* and transgenic hairy roots. **a**) *EXPB1* (tYFPnls reporter construct shown in green), *NPL* (nls-mCherry reporter construct shown in red) and *EXPB1-NPL* coexpression (shown in yellow) expression in wt hairy roots 5 dpi with *M. loti* R7A. **b**) *STF1* (tYFPnls reporter construct shown in green), *NPL* (nls-mCherry reporter construct shown in red) and *STF1-NPL* coexpression (shown in yellow) expression in wt hairy roots 6 dpi with *M. loti* R7A. Scale bar: 100 µm.

**Supplemental Figure S6.**
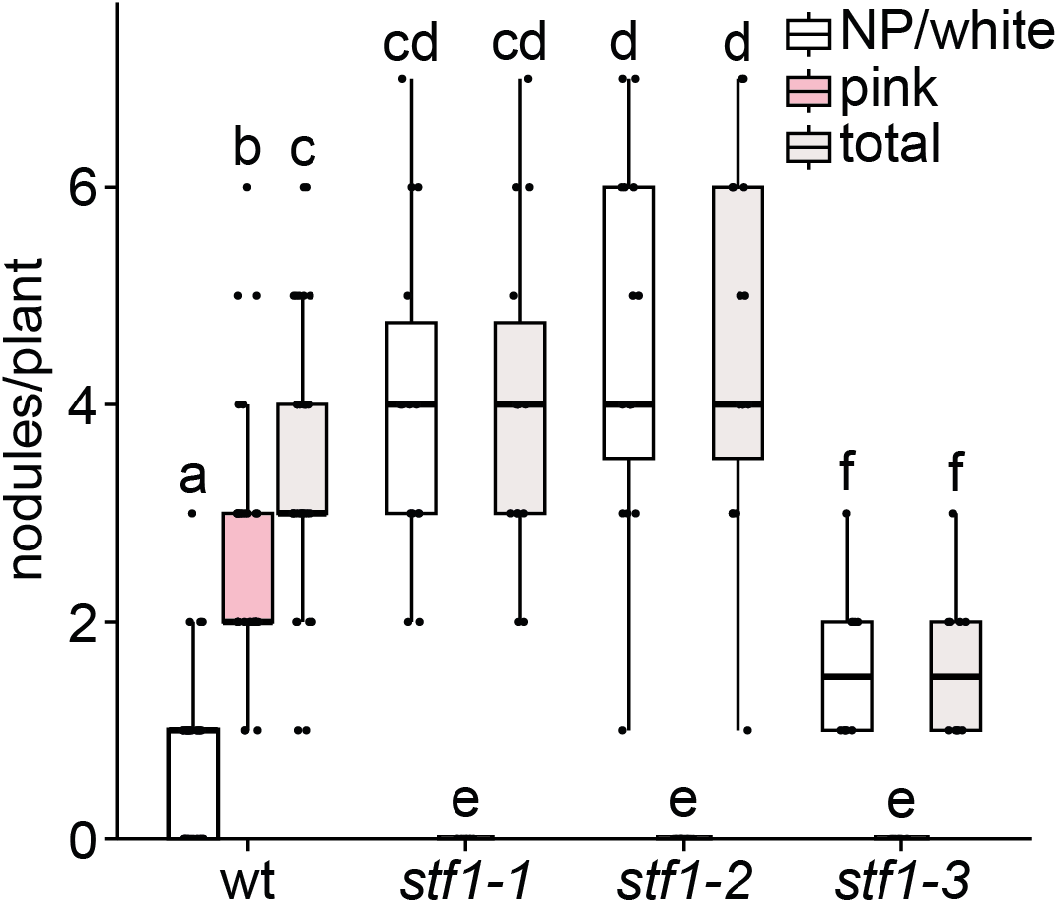
Number of white, pink and total nodules at wild-type (wt), *stf1-1, stf1-2* and *stf1-3* plants 12 dpi with *M. loti* R7A-dsRed (p ≤ 0.05; Scheirer–Ray–Hare with Benjamini-Hochberg-corrected Wilcoxon rank sum test; n ≥ 10 per genotype). Letters indicate significantly different statistical groups.

**Supplemental Figure S7.**
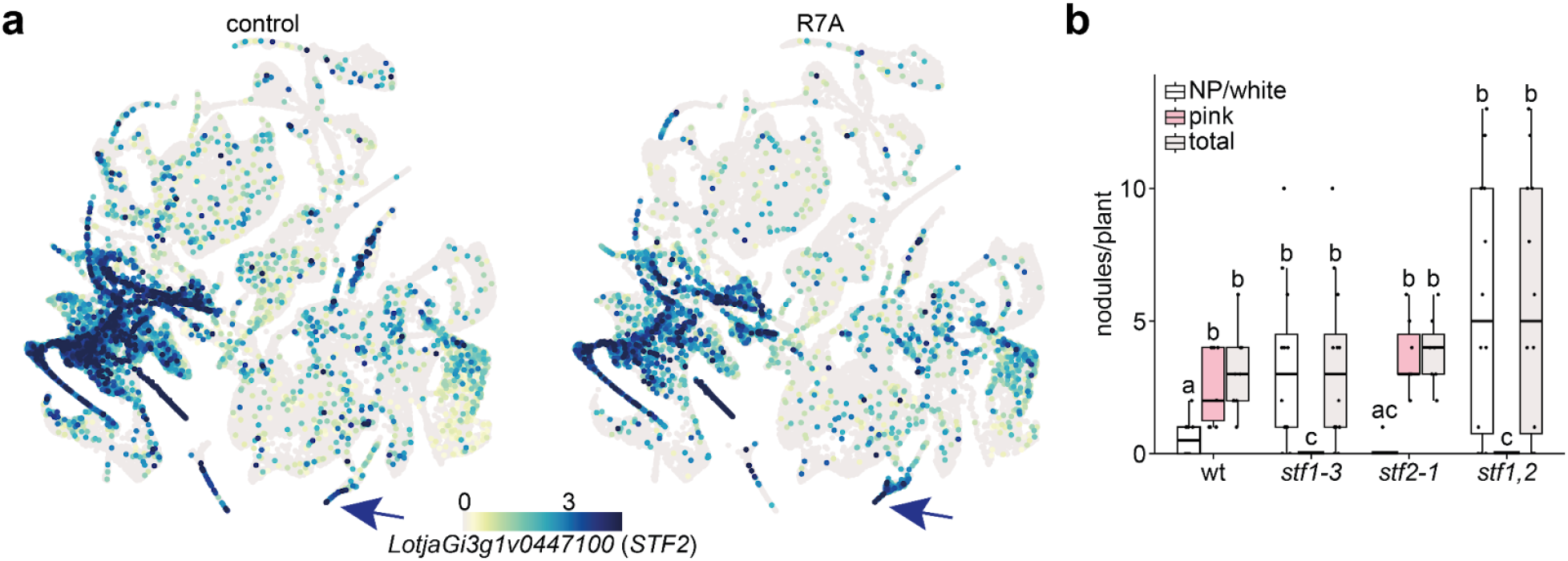
STF2 is not required for successful infection. **a**) Normalised *STF2* expression in all wt cells. Arrows indicate responsive RHs. **b**) Number of white, pink and total nodules at wild-type (wt), *stf1-3, stf2-1* and *stf1,2* plants 42 dpi with *M. loti* R7A (p ≤ 0.05; Scheirer–Ray–Hare with Benjamini-Hochberg-corrected Wilcoxon rank sum test; n ≥ 7 per genotype). Letters indicate significantly different statistical groups.

**Supplemental Figure S8.**
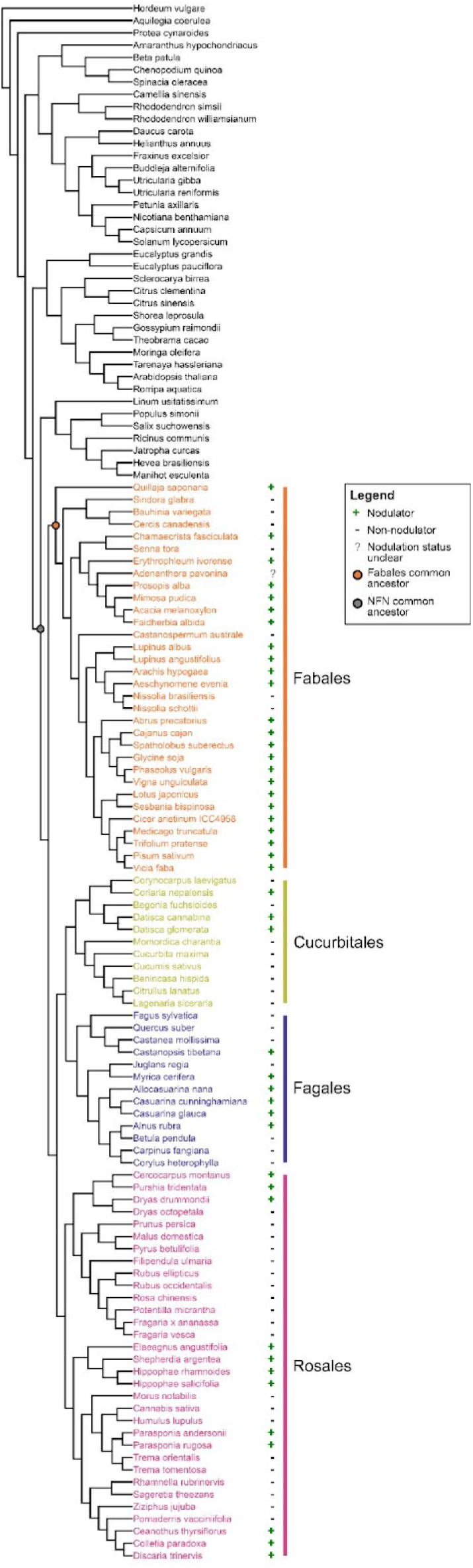
Phylogenetic tree of species used for phylogenomic analyses. Additional information over the species analysed can be found in Source Data “Supp data S5 - Phylogenomics” sheet “genomes”.

**Supplemental Figure S9.**
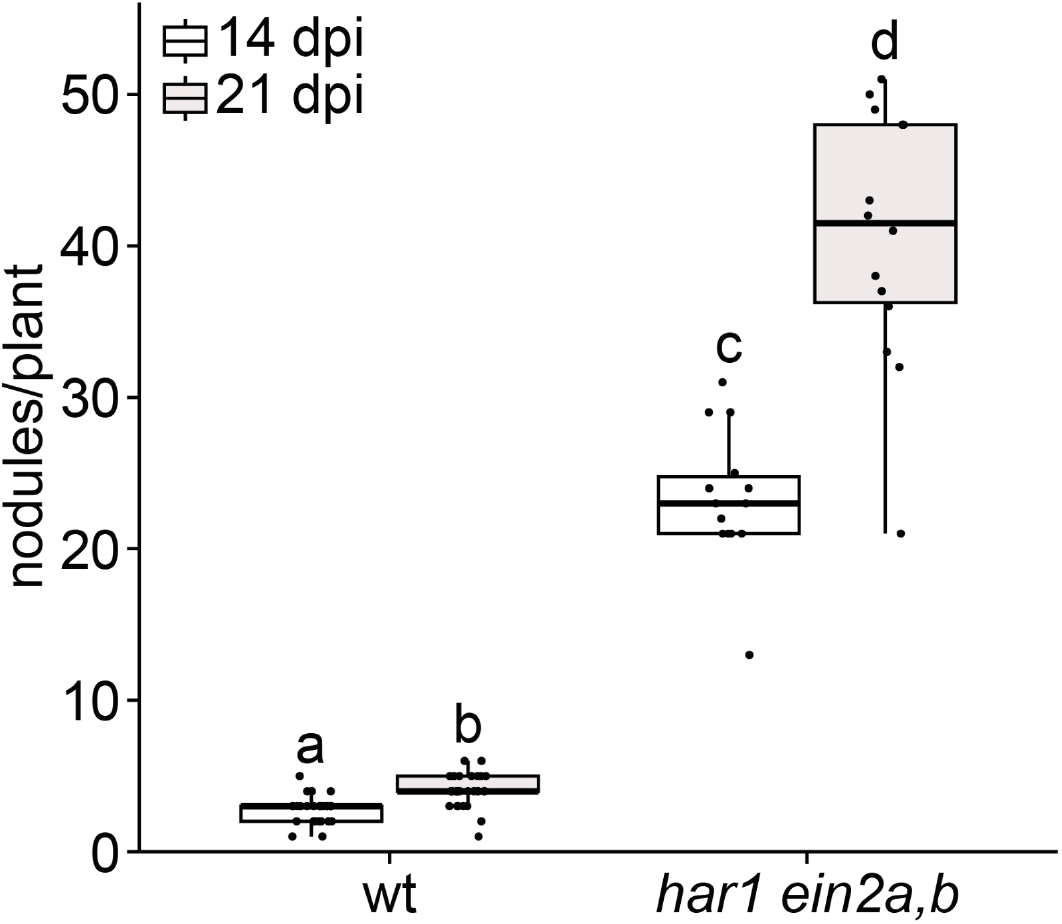
*har1 ein2a,b* displays a hypernodulation phenotype. Number of nodules at wild-type (wt) and *har1 ein2a,b* plants 14 and 21 dpi with *M. loti* R7A-dsRed (p ≤ 0.05; Scheirer–Ray–Hare with Benjamini-Hochberg-corrected Wilcoxon rank sum test; n ≥ 14 per genotype). Letters indicate significantly different statistical groups. Raw data and statistical analysis for the displayed experiment are available in Source Data “Supp data S3 - nodule and IT quantifications”.

**Supplemental Figure S10.**
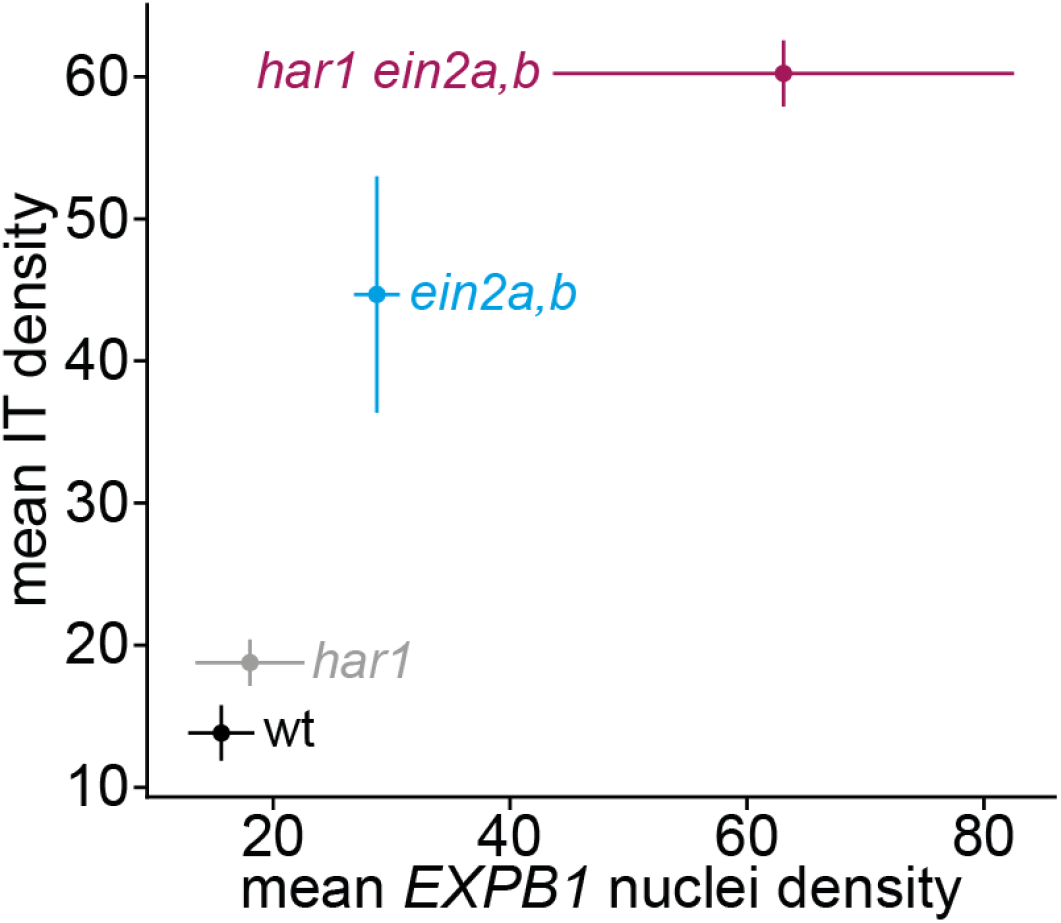
Relationship between susceptible RH abundance and IT density across genotypes. Mean EXPB1::tYFPnls-positive RH density (x-axis) and mean IT density (y-axis) were quantified independently in wild-type (wt), *har1, ein2a,b* and *har1 ein2a,b* plants. Each point represents the genotype mean ± s.e.m. for EXPB1-positive RH density (horizontal error bars) and IT density (vertical error bars). Genotypes with higher EXPB1-positive RH density also exhibited higher IT density, with both measures increasing from wt through *har1* and *ein2a,b* to *har1 ein2a,b* (Spearman ρ = 1.0 across four genotype means; Source Data “Supp data S8 - Spearman”).

**Supplemental Figure S11.**
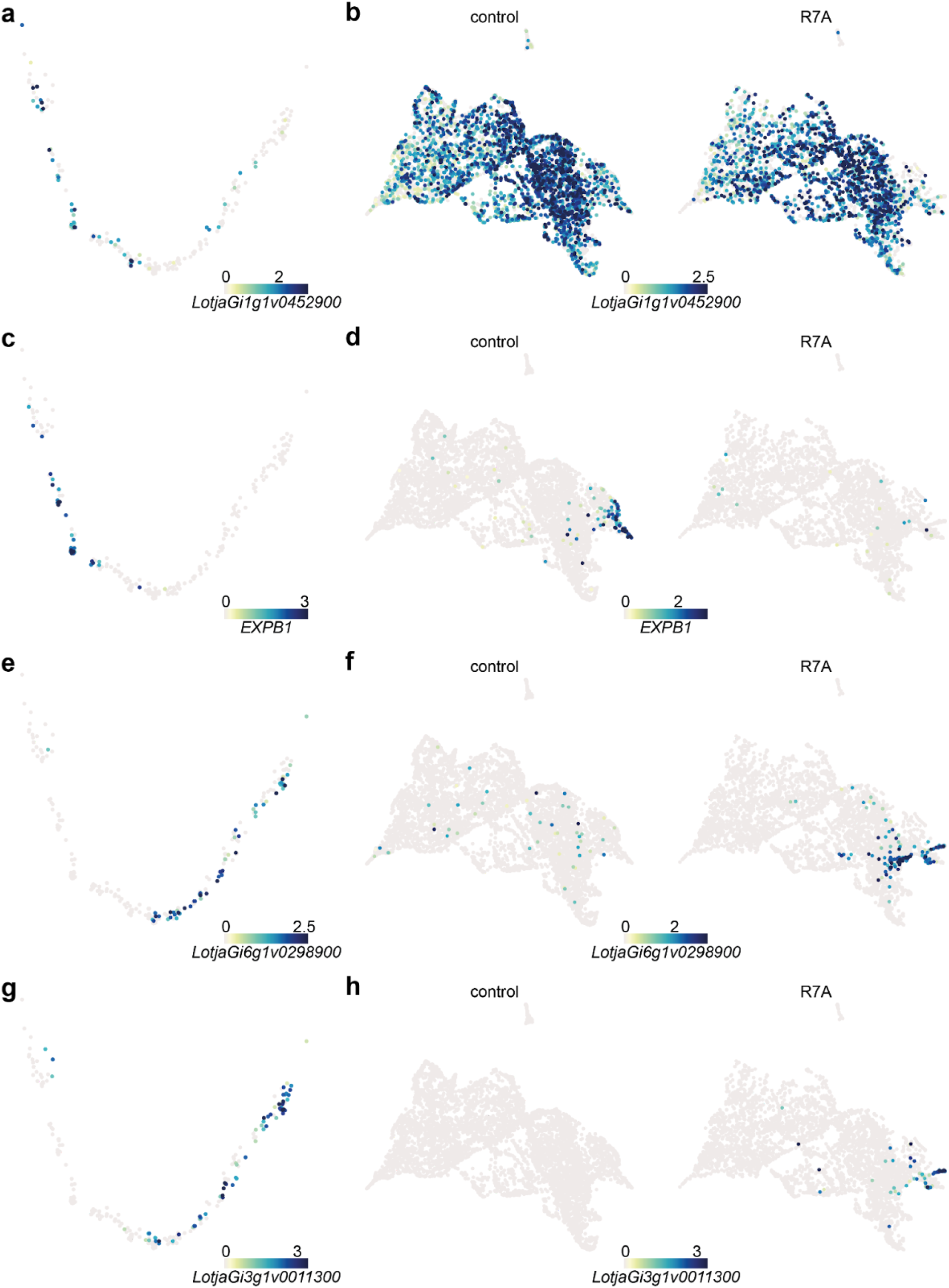
Normalised pseudotime cluster marker gene expression in wt and *har1 ein2a,b* RHs. Normalised gene expression in cluster 21 RHs (**a, c, e** and **g**) and all RHs (**b, d, f** and **h**).

